# On the role of L-type Ca^2+^ and BK channels in a biophysical model of cartwheel interneurons

**DOI:** 10.1101/2025.08.01.668076

**Authors:** Matteo Martin, Jonathan E. Rubin, Morten Gram Pedersen

## Abstract

Cartwheel interneurons (CWCs) in the auditory system exhibit a range of activity patterns relevant to auditory function and pathologies. Although experiments have shown how these patterns can vary across individual neurons and can change under pharmacological manipulations, the field has lacked a computational framework in which to explore the contributions of particular currents to these observations and to generate new predictions about the effects of manipulations on CWCs. In this work, we address this deficiency by presenting a conductance-based CWC computational model. This model captures the diversity of CWC activity patterns observed experimentally and suggests parameter changes that may underlie differences across cells. Bifurcation analysis of this model provides an explanation of how distinct dynamic mechanisms contribute to these differences, while direct simulations suggest how cells with different baseline dynamics will respond to variations in certain experimentally-accessible potassium and calcium channel conductances. In addition to the full model that we introduce, we present a reduced model that preserves CWC dynamic regimes. We classify the reduced model variables in terms of distinct dynamic timescales and show that the key transitions in dynamic patterns can be explained based on equilibria of the averaged dynamics of the slowest model variables, in a regime where the faster model variables exhibit oscillations. Overall, this study predicts how changes in parameters will influence CWC behavior, suggests how bifurcations contribute to changes in CWC dynamics, and provides a theoretical foundation that supports our simulation findings.

**Author summary:** Cartwheel interneurons (CWCs) are the most common class of inhibitory interneurons in an auditory brainstem region involved in sound localization and are believed to be important for auditory processing and pathologies. Distinct patterns of CWC activity have been observed experimentally in a variety of conditions. In this work, we present two novel computational models that simulate the factors contributing to CWC dynamics. By harnessing this framework, we are able to reveal the contributions of key ion currents to modulating CWC activity. Indeed, we find that the factors present in CWC neurons can produce a complicated dynamic landscape, with a wide range of output patterns possible as the relative strengths of these factors are varied. Overall, our models represent useful tools for understanding experimental results and generating new predictions about CWC behavior. In particular, in the more reduced of the two models, we can perform mathematical analysis to make more detailed predictions about the effects of current modulation on whether CWC neurons will exhibit regular spiking or more complex forms of outputs.

## Introduction

Cartwheel cells (CWC) are cross-species [1–6], glycinergic interneurons located in the dorsal cochlear nucleus (DCN). As the predominant form of inhibitory neuron in the DCN [7], they regulate the electrical activity of fusiform neurons [8] and thus they can play a role in auditory processing as well as in pathologies such as tinnitus [9–11]. To understand auditory function and dysfunction, it is therefore crucial to study the mechanisms driving and modulating CWC electrical activity.

CWCs exhibit diverse activity patterns classified as spiking, bursting, and complex spiking [12]. Complex spikes include both large amplitude action potentials (LAOs) and smaller amplitude oscillations (SAOs), which Kim et al. call spikelets [12]. While certain forms of bursting can also feature SAOs, complex spiking differs from neuronal bursting [13–15] in that it lacks the prolonged epochs of quiescence between groups of spikes that occur in burst patterns. Functionally speaking, complex spiking may play a crucial role in microcircuit network communication by influencing the susceptibility of neighboring neurons to Ca^2+^-dependent plasticity [16]. Experimental studies have distinguished spiking from complex spiking CWCs based on the oscillation patterns that they display at the onset of activity as an applied current is varied. Both CWC types, however, can exhibit forms of complex spiking in response to sufficient increases in depolarizing applied current. Moreover, treatment with nifedipine (an inhibitor of L-type Ca^2+^ channels) can upregulate the likelihood of activating regular spiking, while treatment with iberiotoxin (an inhibitor of big-conductance, Ca^2+^-dependent K^+^ (BK) channels) can downregulate the occurrence of regular spiking, in both classes of CWCs [12, 17].

In this work, we introduce a novel eleven-dimensional, conductance-based ordinary differential equation model [18] for CWCs, which we use to analyze the dynamic mechanisms underlying these observations about CWC dynamics. Specifically, we first use one-parameter bifurcation analysis to show how alterations in the balance of calcium and potassium currents in CWCs can lead to the observed differences in spiker and complex spiker dynamics at activity onset. We next show that the model can reproduce experimental results where BK or L-type channels were blocked by iberiotoxin or nifedipine, respectively, and we then explore effects of more modest reductions, as well as increases, of BK and L-type Ca^2+^ conductances. With a model, we can consider both the pharmacological lesions performed experimentally as well as more subtle changes in channel activity, such as the effects of lower drug doses [19, 20], neuromodulation [21], or gain-/loss-of-function mutations, transcription or trafficking defects, and other consequences of channelopathies [22]. To this end, we create heatmaps (cf. [23]) illustrating how a morphological index, defined as an extension of the firing number [24], changes when the applied current (*I*_*App*_) and the BK or L-type Ca^2+^ conductance (*g*_*BK*_ or *g*_*CaL*_) are varied. We further investigate model responses to simulated variations in BK and L-type conductances by introducing a model reduction and applying averaging theory [25, 26]. The results of this investigation demonstrate a relation between the spiking patterns observed under parameter variation in simulations and the stability properties of an equilibrium point of a superslow component of the model system. Overall, our models can be used to predict how parameter changes will affect CWC behavior and to identify ways to alter CWC dynamics, and they provide a theoretical underpinning that substantiates our simulation results.

## Results

### Two classes of CWC conductance-based model dynamics

We developed a novel conductance-based CWC interneuron model, based on a variety of experimental observations about these cells (see *Methods*, equations (1)). We used two different parameter sets for our model to exemplify two different classes of CWC interneurons, spikers and complex spikers, defined based on the voltage dynamics they exhibit at the onset of activity as an applied current, *I*_*App*_, is increased from hyperpolarizing levels. Specifically, the spiker model compared to the complex spiker has lower whole-cell conductances of delayed-rectifier K^+^ (*g*_*K*_) and L-type Ca^2+^ (*g*_*CaL*_) channels, but higher persistent Na^+^ (*g*_*NaP*_) and Ca^2+^-dependent K^+^ (*g*_*K*(*Ca*)_) conductances.

For these combinations of parameters, the two classes of cells produce progressions of activity patterns that agree with experimental observations as *I*_*App*_ is increased [12]. For low values of *I*_*App*_, complex spikers and spikers exhibit pseudo-plateau bursting and regular spiking, respectively, as shown in Fig 1A. For larger *I*_*App*_, the two types of interneurons both engage in regular spiking (Fig 1B), and finally, for high enough *I*_*App*_, the interneurons generate complex spiking activity (Fig 1C).

**Fig 1.**
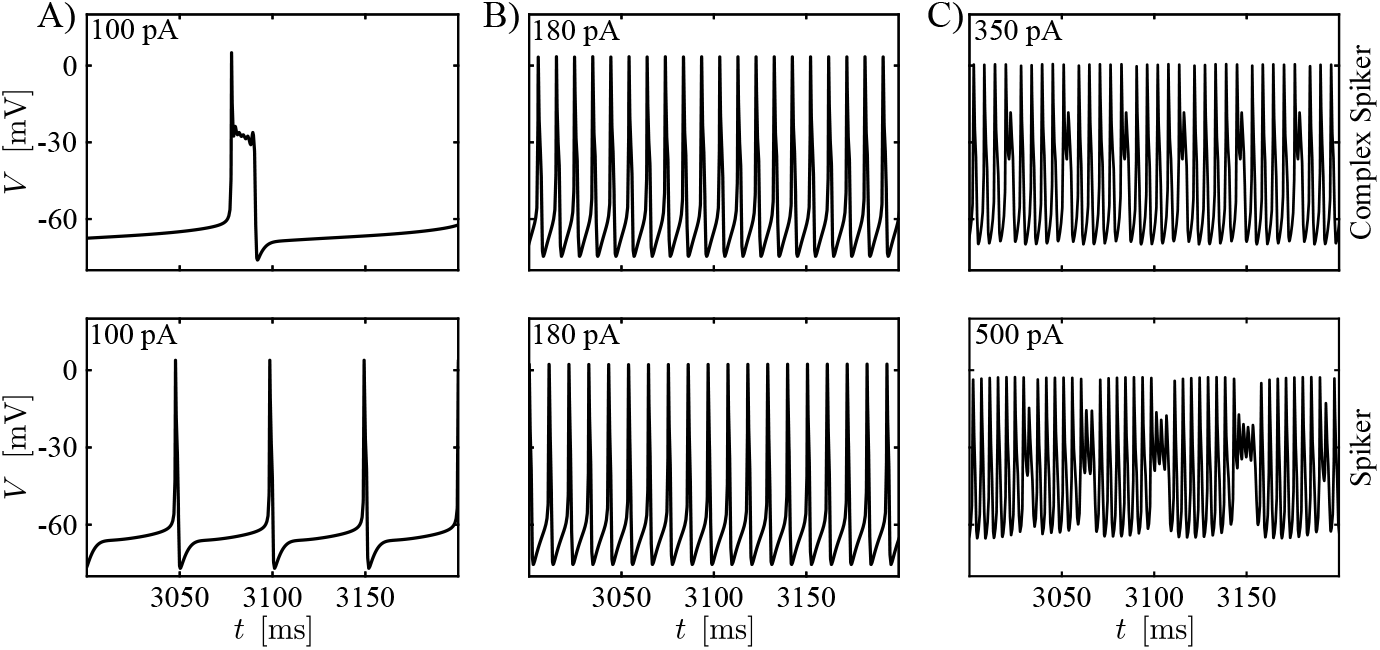
CWC interneuron model time series. (A) At low *I*_*App*_, the complex spiker model exhibits pseudo-plateau bursting (top), whereas the spiker model displays spiking. (B) Both the complex spiker (top) and the spiker models show regular spiking for larger *I*_*App*_. (C) Both models produce complex spiking for still greater *I*_*App*_, although larger values are needed for the spiker (bottom) than for the complex spiker. In all panels, the *I*_*App*_ value used appears in the upper left.

To characterize the model dynamics and the difference between the two versions of the model more thoroughly, we computed bifurcation diagrams (BD) for both the complex spiker and spiker models, with *I*_*App*_ as the bifurcation parameter (Figs 2A and 2B). Each bifurcation diagam in Fig 2 shows a variety of labeled bifurcation events. We denote the *I*_*App*_ value where a bifurcation B occurs by *I*_*App*,B_, where B can denote any bifurcation type. Along with each *I*_*App*_-BD, we include a bar above the main plot showing how the firing number Φ [24] changes with *I*_*App*_. This index is defined as Φ = *L/*(*L* + *s*) where *L* is the number of large-amplitude oscillations and *s* is the number of small-amplitude oscillations occurring per oscillation cycle within the system’s stable oscillation pattern. It takes the value 1 for regular, large amplitude spiking, decreases as small amplitude oscillations become more prevalent, and is equal to 0 for tunings in which the neuron is silent. More details are provided in the Firing number section of *Methods*. The bars on the right of the *I*_*App*_-BDs show the mapping between shades of grey and values of Φ. In particular, black and white correspond to a zero and unitary Φ, respectively.

**Fig 2.**
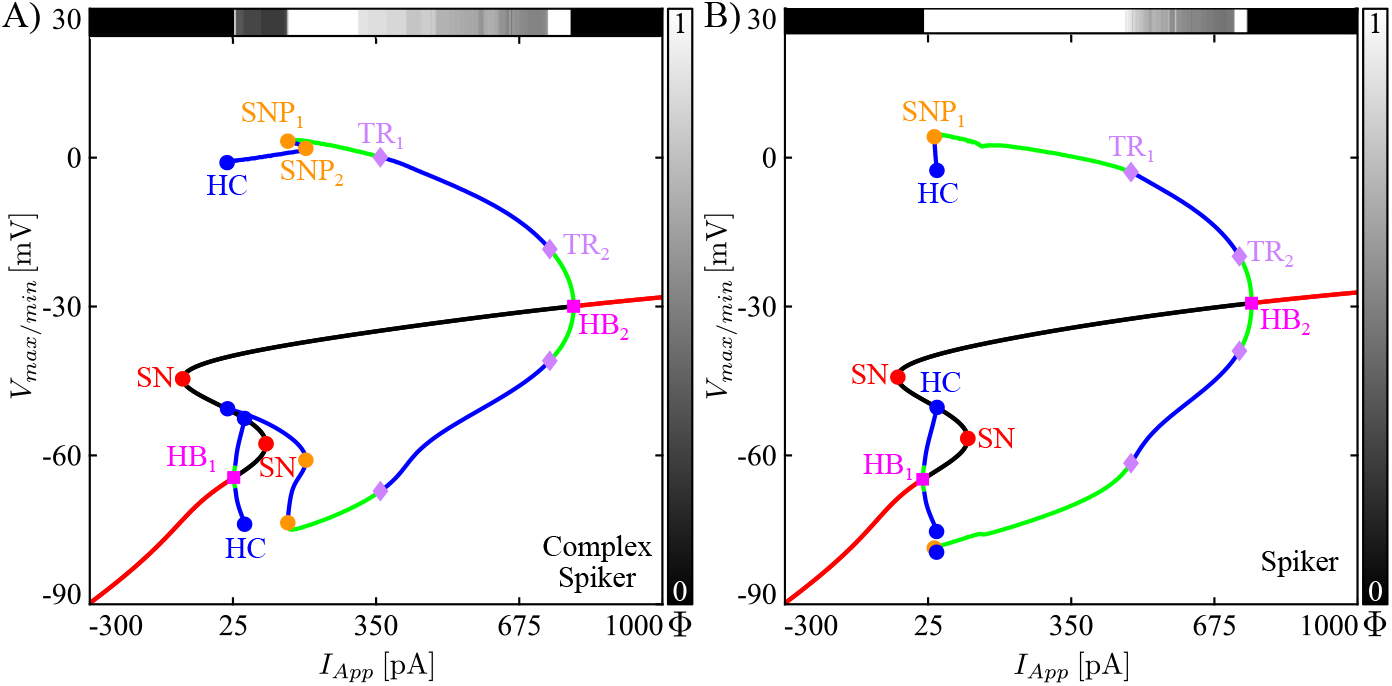
Bifurcation diagrams for the CWC model. Bifurcation diagrams with respect to *I*_*App*_ for two parameter sets corresponding to complex spikers (A) and spikers (B). Labeled geometric symbols denote bifurcations: The orange (red) dots are associated with saddle-node bifurcations of periodic orbits (SNP) (of critical points (SN)), the blue dots with homoclinic bifurcations (HC), the magenta squares with Andronov-Hopf bifurcations (HB), the violet diamonds with torus bifurcations (TR). The red/black continuous curve indicates *V* values for stable/unstable critical points. The green/blue curves represent the maximum and the minimum voltages along stable/unstable periodic orbits. The horizontal bar along the top of the BD shows how the firing number Φ changes along the system’s attracting solution with variations in *I*_*App*_. The greyscale coding used for this bar is indicated on the right (white: Φ = 1; black: Φ = 0). In particular, white intervals in the top bar correspond to regular spiking regimes.

The branch of equilibria in each of the BDs in Fig 2 has the typical S shape with folds at saddle-node (SN) bifurcations, as seen for many models of cellular transmembrane electrical activity. We observe that both model neurons are silent if a hyperpolarizing (negative) current is applied. This situation persists as current is increased through zero until it becomes depolarizing and reaches 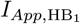, which is equal to 12 (27) pA in the spiker (complex spiker) model, where a supercritical Andronov-Hopf bifurcation (HB_1_) occurs. At this critical value, the stable equilibrium point loses its stability, and the two models start to behave differently. In both cases, although HB_1_ is supercritical and hence gives rise to a family of periodic solutions that are initially stable, these solutions destabilize almost immediately and remain unstable throughout the remainder of their extent, up through termination in a homoclinic bifurcation (HC).

For the complex spiker, over a significant interval of *I*_*App*_ values above 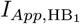, no stable solution is evident in the bifurcation diagram (Fig 2A). Here, the complex spiker exhibits a form of activity that combines LAOs and SAOs (Fig 1A), known as pseudo-plateau bursting, with small Φ. With additional increases in *I*_*App*_, the complex spiker eventually undergoes a saddle-node of periodic orbits bifurcation (SNP_1_) and switches to regular spiking with Φ = 1 (green curves between SNP_1_ and TR_1_, Fig 2A). For the spiker (Fig 2B), on the other hand, at an *I*_*App*_ value just slightly above 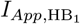, a saddle-node of periodic orbits bifurcation (SNP_1_) occurs, which gives rise to a stable family of large amplitude, periodic spiking solutions (green curves between SNP_1_ and the torus bifurcation TR_1_). Hence, the spiker exhibits an almost-immediate transition from quiescence to regular spiking, with Φ = 1.

Overall, the key features that distinguish the complex spiker from the spiker in this model are: the added complexity of the leftmost part of the branch of periodic orbits in the complex spiker BD for *I*_*App*_ between 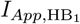 and 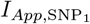, the larger value of 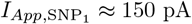 where the onset of regular spiking occurs for the complex spiker, and the significantly wider interval of *I*_*App*_ values yielding regular spiking for the spiker case.

If we increase *I*_*App*_ far enough above 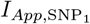, both models’ dynamics switch to complex spiking. We notice that, contrary to the previous cases, where the transitions between different states were directly associated with bifurcations in the *I*_*App*_-BD, the transition from spiking to complex spiking is not. Specifically, the transition occurs before the critical value 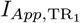, which is equal to 485 and 359 pA in the spiker and complex spiker models, respectively. Indeed, for the complex spiker, as *I*_*App*_ increases from 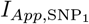 toward 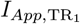, the firing number Φ starts to decrease before 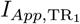 is reached, reflecting the change in the signature of the model’s dynamics. Various forms of complex spiking activity persist in both models until the critical value of 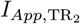, which is equal to 732 and 744 pA in the spiker and complex spiker models, respectively, is reached. For *I*_*App*_ above 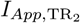, the models switch to stable, high-frequency but low-amplitude, regular spiking. Finally, both models enter a state of depolarization block with a stable equilibrium once *I*_*App*_ exceeds the Andoronov-Hopf bifurcation HB_2_ at 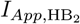, equal to 798 (759) pA in the complex spiker (spiker) model.

In summary, raising *g*_*K*_ and *g*_*CaL*_ and lowering *g*_*NaP*_ and *g*_*K*(*Ca*)_ alter spiker dynamics in two main ways. First, these changes interrupt the direct transition from quiescence to regular spiking via the introduction of a significant window of *I*_*App*_ values over which pseudo-plateau bursting occurs and an increase in the *I*_*App*_ value needed to achieve regular spiking. Second, they shrink the interval of *I*_*App*_ values over which regular spiking occurs, allowing complex spiking to kick in at relatively much lower *I*_*App*_.

### Inhibition of BK channels promotes complex spiking and bursting in CWCs

Iberiotoxin, a potent big-conductance K^+^ (BK) channel blocker, diminishes the likelihood of observing regular spiking in cartwheel interneurons and promotes the emergence of bursting or complex spiking phenomena [12, 17]. Our model can reproduce these results and allows us to explore and predict effects of gradual reductions or increases in the BK channel conductance, *g*_*BK*_, across a range of *I*_*App*_ values.

Figs 3A and 4A display results of systematic variation of *g*_*BK*_ and *I*_*App*_ for our two baseline model tunings in terms of heatmaps of the firing number Φ. In these figures, the baseline value of *g*_*BK*_ is indicated with a black, dashed horizontal line. Before exploring less extreme modifications of the BK conductance, we simulate the iberiotoxin experiments, assuming that the drug blocks all of the BK channels in the cell. The sample trajectories for the complex spiker presented in Fig 3B show that for *I*_*App*_ equal to 180 pA, the model under control conditions (*g*_*BK*_ = 80 nS) produces regular spiking, which is converted into pseudo-plateau bursting when iberiotoxin is administered (*g*_*BK*_ = 0 nS). For a stimulating current of 350 pA, the complex spiker produces complex spiking, which persists with a smaller Φ value under iberiotoxin administration (*g*_*BK*_ = 0 nS). Fig 4B shows the behavior of a spiker interneuron under control (*g*_*BK*_ = 80 nS) and iberiotixon-treated (*g*_*BK*_ = 0 nS) conditions. For *I*_*App*_ equal to 180 pA or 350 pA, the spiker interneuron exhibits regular spiking, which is transformed into pseudo-plateau bursting or complex spiking, respectively, when iberiotoxin is administered. Thus, both models exhibit similar responses to iberiotoxin, despite their differences in baseline dynamics.

**Fig 3.**
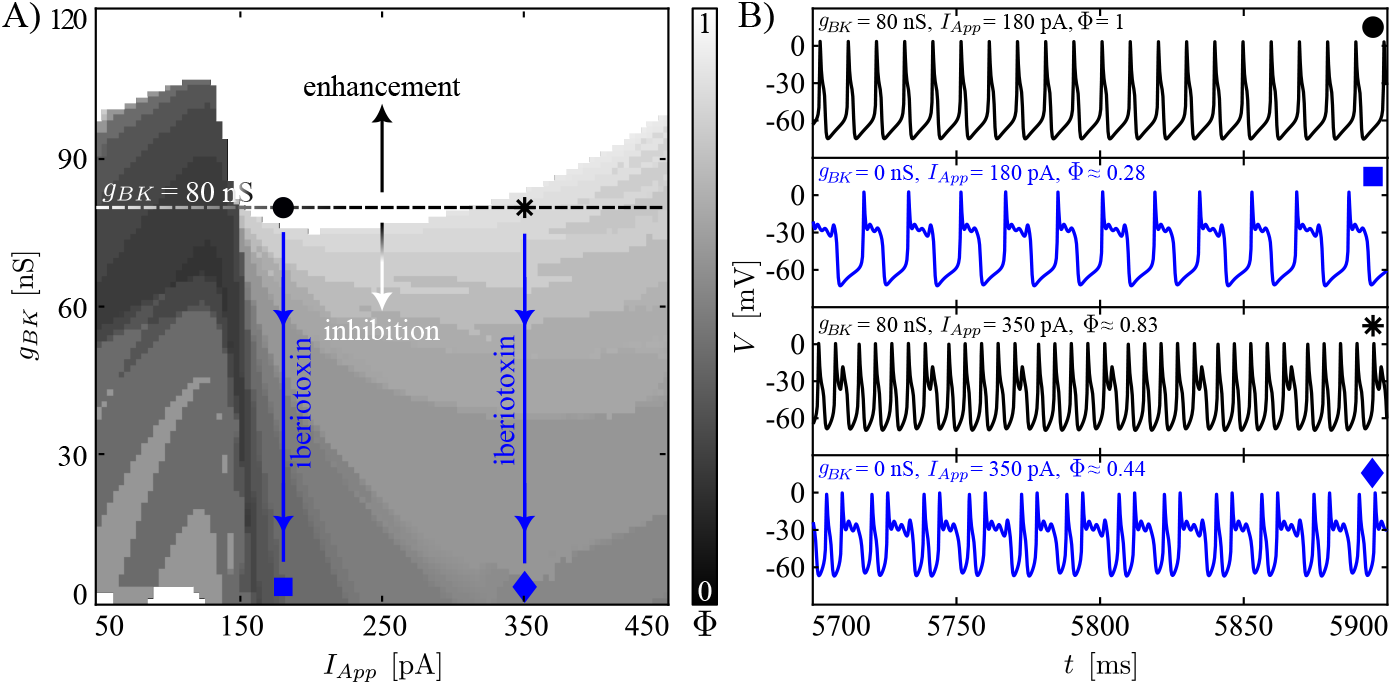
Responses of complex spikers to variation of *g*_*BK*_ and *I*_*App*_ and to iberiotoxin adminstration. (A) Heatmap for the index Φ with respect to parameters (*I*_*App*_, *g*_*BK*_). The vertical bar on the right of the plot corresponds to firing number Φ (cf. Fig 2). The horizontal black dashed line represents the baseline value of *g*_*BK*_ (80 nS) used in the model. The regular spiking (white) in this type of interneuron is highly sensitive to small reductions of *g*_*BK*_ because the baseline dynamics is set close to the border of the regular spiking domain. The vertical, blue lines represent effects of iberiotoxin administration. Each connects two symbols, the circle/asterisk and the square/diamond, respectively. (B) Model behavior at the parameter values indicated by the four markers presented in (A). Blockade of *g*_*BK*_ converts regular spiking into bursting, while complex spiking persists with more SAOs relative to LAOs (smaller Φ).

**Fig 4.**
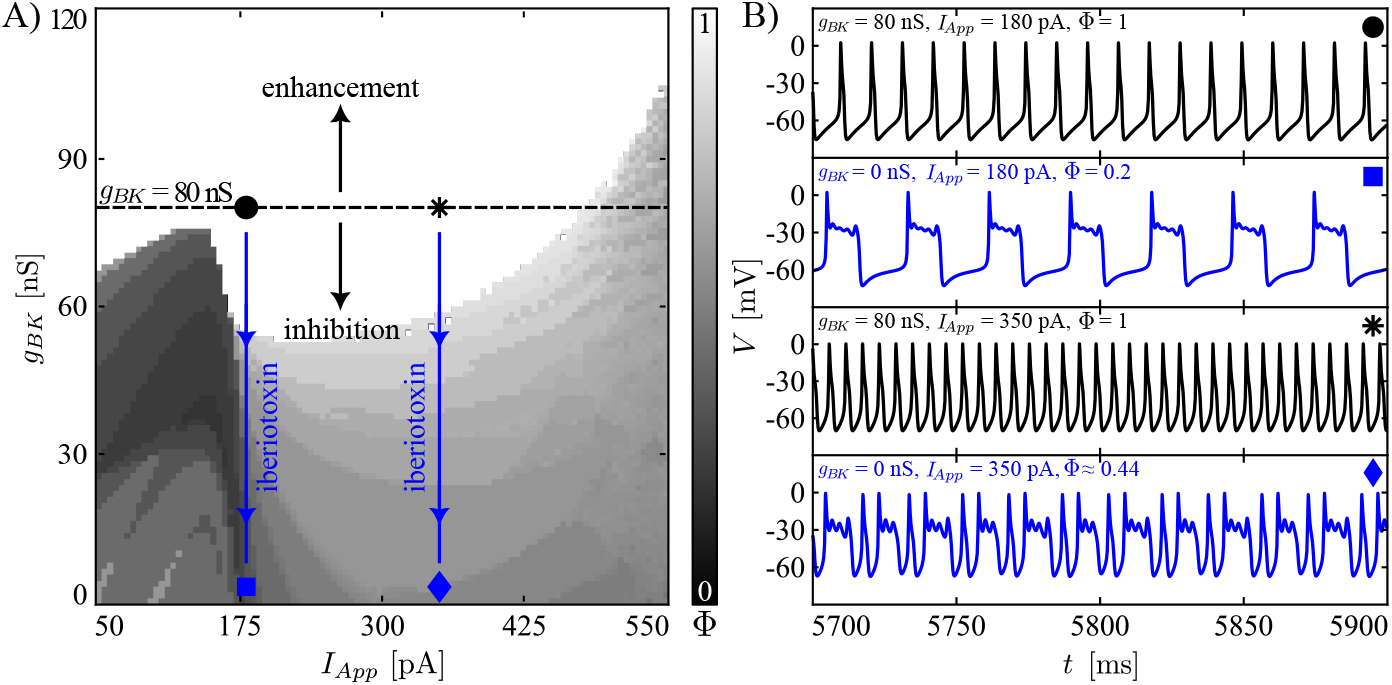
Responses of spikers to variation of *g*_*BK*_ and *I*_*App*_, and to iberiotoxin adminstration. Notation follows that of Fig 3. (A) Heatmap of the firing number Φ under variation of the two parameters *g*_*BK*_ and *I*_*App*_. (B) Model dynamics at the different points in parameter space are indicated by markers of various shapes. Simulated iberiotoxin administration converts regular spiking to pseudo-plateau bursting (*I*_*App*_ = 180 pA) or complex spiking (*I*_*App*_ = 350 pA).

Next, we consider more gradual reductions of *g*_*BK*_ (regions of the heatmaps below the black, dashed horizontal line). In the heatmaps (Figs 3A and 4A), the color-coding of Φ corresponds to that in Fig 2. In both cases, we see that starting from a regular spiking regime achieved by tuning of *I*_*App*_, reductions of *g*_*BK*_ cause decreases in Φ, representing a larger number of SAOs and/or a lower number of LAOs in the model’s periodic activity patterns. The same trend, with fewer LAOs and more SAOs as *g*_*BK*_ decreases, recurs for the complex spiker case when *I*_*App*_ is increased to 350 pA, corresponding to what is initially a complex spiking regime (Fig 3B). For the complex spiker, the region of regular spiking with *g*_*BK*_ equal to 80 nS is especially fragile, with small reductions in *g*_*BK*_ eliminating regular spiking (Fig 3A). This outcome arises because the complex spiker model, due to the chosen parameter settings, lies close to the border of the regular spiking domain. Another small window with Φ = 1 emerges for the complex spiker with *I*_*App*_ small and *g*_*BK*_ close to 0 nS. However, unitary Φ values in this region do not correspond to regular spiking but rather to plateau-bursting with no SAOs during the active phase, i.e., very broad action potentials, each consisting of a spike and a depolarized plateau, separated by prolonged pauses. For the spiker case (Fig 4A), the regular spiking area around the *g*_*BK*_ baseline value (80 nS) is wider and more robust than that of the complex spiker. In this regime, a reduction of *g*_*BK*_ of at least 35% is needed to abolish regular spiking.

In contrast to the previous observations, when *g*_*BK*_ increases (regions of the heatmaps above the black, dashed horizontal lines), in both the complex spiker and the spiker, the range of *I*_*App*_ yielding regular spiking widens. In general, we note that although the switch between the complex spiker and spiker models involves changes in several conductances, the two models produce qualitatively similar heatmaps with respect to (*I*_*App*_, *g*_*BK*_), with a quantitative shift in the *g*_*BK*_ values associated with similar dynamics across the two models.

From the biological perspective, the BK channels activate along with P/Q-type Ca^2+^ currents and act on a depolarized transmembrane potential, providing a hyperpolarizing effect. When the BK conductance is set to 0 nS, one of the forces contributing to repolarization of the membrane potential towards the resting state is no longer present. For this reason, the membrane potential evolves around a depolarized plateau, where small fluctuations may occur. Both the regular spiking and the complex spiking cases (with few SAOs) start to show longer depolarized plateaus with a higher number of SAOs superimposed.

### Inhibition of the L-Type Ca^2+^ channel enhances regular spiking

The L-type Ca^2+^ channel current can be suppressed by the pharmacological agent nifedipine. Experimental application of this blocker abolished complex spiking activity in most CWC interneurons [12]. Our model reproduces this result and allows us to explore and predict the effects of varying degrees of reduction and increase of the L-type Ca^2+^ channel maximal conductance *g*_*CaL*_ across a range of *I*_*App*_ values.

We once again use heatmaps to show how changes in parameters, in this case (*I*_*App*_, *g*_*CaL*_), yield variations in the firing number Φ (Fig 5A and Fig 6A) from the baseline condition (represented using an horizontal black dashed line). Analogous to the iberiotoxin case, the administration of nifedipine corresponds to a vertical jump from the maximum value of L-type Ca^2+^ conductance (control case), 19 (23) nS for the spiker (complex spiker) interneuron model, towards a value of 0 nS (nifedipine-treated conditions). Figs 5B and 6B show sample trajectories depicting how nifedipine alters the electrical activity in the complex spiker and spiker models, consistently producing regular action potential firing at the expense of more complex patterns.

**Fig 5.**
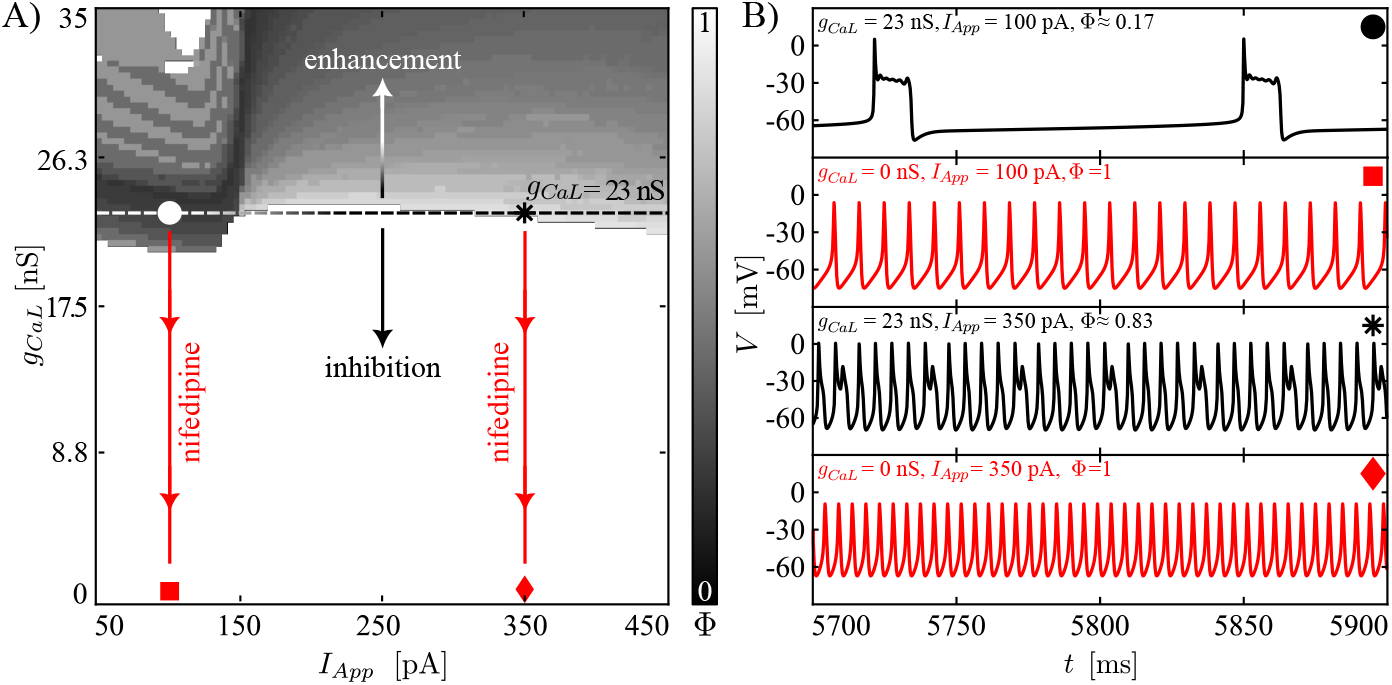
Responses of complex spikers to variation of *g*_*CaL*_, *I*_*App*_, and to nifedipine adminstration. The content of the current figure is organized as in Fig 3. (A) Heatmap showing how the firing number Φ changes as *I*_*App*_ and *g*_*CaL*_ are varied. (B) Time series associated with the markers presented in panel (A), corresponding to, from top to bottom, control conditions with relatively low *I*_*App*_, nifedipine administration with relatively low *I*_*App*_, control conditions with relatively high *I*_*App*_, and nifedipine administration with relatively high *I*_*App*_.

**Fig 6.**
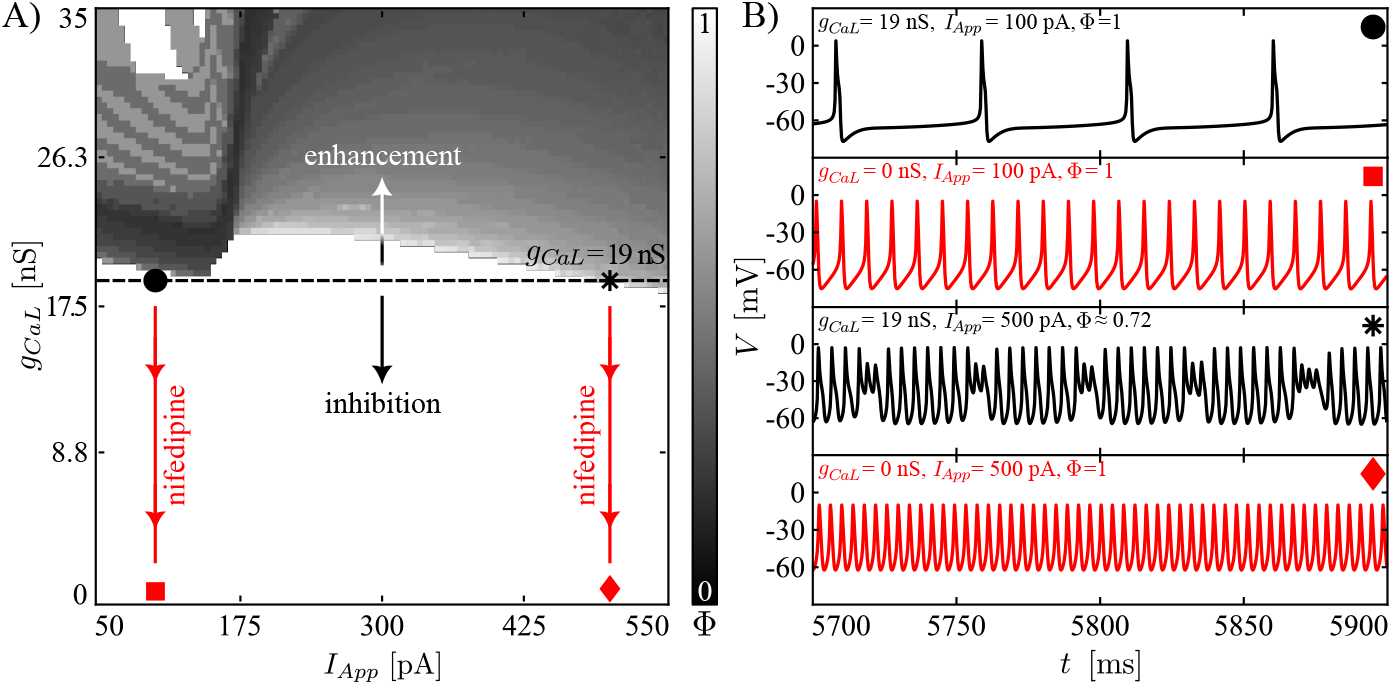
Responses of spikers to variation of *g*_*CaL*_, *I*_*App*_, and to nifedipine adminstration. The content of the current figure is organized as in Fig 3. (A) Heatmap showing how the firing number Φ changes as *I*_*App*_ and *g*_*CaL*_ are varied. (B) Time series associated with the markers presented in panel A, corresponding to, from top to bottom, control conditions with relatively low *I*_*App*_, nifedipine administration with relatively low *I*_*App*_, control conditions with relatively high *I*_*App*_, and nifedipine administration with relatively high *I*_*App*_.

In the model of a complex spiker, with default *g*_*CaL*_ = 23 nS and low *I*_*App*_, Φ is small because the complex spiker exhibits pseudo-plateau bursting (Fig 5A). The two Φ *<* 1 regions (grey) at the control level of *g*_*CaL*_ (for *I*_*App*_ between 50 and 150 pA or for *I*_*App*_ *>* 300 pA) are highly sensitive to small changes in the L-type Ca^2+^ conductance for the chosen parameters, and in both regions regular spiking activity takes over after small reductions in *g*_*CaL*_, exemplified by the simulations of nifedipine administration at *I*_*App*_ equal to 100 or 350 pA (Fig 5B).

Increasing the maximal conductance of *g*_*CaL*_ has instead an opposite effect. In fact, the region of regular spiking activity at baseline *g*_*CaL*_ disappears in response to a small increase in *g*_*CaL*_, while bursting and complex spiking regions persist and expand as *g*_*CaL*_ increases. In agreement with the results presented for the (*I*_*App*_, *g*_*BK*_) heatmap, for *I*_*App*_ between 50 and 150 pA, as *g*_*CaL*_ increases the depolarized potentials arising during bursts stabilize and lengthen. When *g*_*CaL*_ is near 35 nS, this effect yields Φ = 1; that is, the activity pattern in this part of parameter space becomes plateau bursting, such that each cycle features a long active phase with no SAOs superimposed.

Fig 6A presents analogous results for a spiker interneuron. The heatmap is predominantly white (Φ = 1); as observed for the complex spiker, the complex spiking activities that arise for large *I*_*App*_ are completely abolished after a small drop in *g*_*CaL*_. Fig 6B shows some trajectories depicting how the neuron behaves as the *g*_*CaL*_ conductance is changed between control and nifedipine-treated conditions for *I*_*App*_ ∈ {100, 500} pA, including the loss of complex spiking for *I*_*App*_ = 500 pA. In the spiker model, a small increase in the maximal conductance of the L-type Ca^2+^ channel maintains complex spiking for large *I*_*App*_ and leads to the generation of bursting phenomena for low values of *I*_*App*_. As for the complex spiker scenario, when *g*_*CaL*_ is close to 35 nS and the value of *I*_*App*_ is between 50 and 150 pA, the model exhibits plateau bursting and the heatmap features unitary Φ.

L-type Ca^2+^ channels activate at a lower voltage compared to the P/Q-type Ca^2+^ channels. They depolarize the cell membrane potential as a result of Ca^2+^ influx. When they are blocked through inhibitors such as nifedipine, the depolarizing forces on the cell membrane are weaker. It is therefore, at first glance, surprising that the cell exhibiting regular spiking becomes more active when inward L-type Ca^2+^ currents are blocked (Fig 6B, upper two traces) [12]. We use our model to explain this behavior as follows. First, the resulting reduction of depolarization yields lower action potential amplitude and consequently less activation of delayed-rectifier K^+^ currents. Second, in the presence of L-type Ca^2+^ channel inhibitors there is less Ca^2+^ influx into the cell, and as a result fewer SK channels activate. The overall reduction in outward K^+^ current causes less afterhypolarization after each spike and hence allows faster regular spiking (Fig 6B, top two traces). The switch from complex spiking or pseudo-plateau bursting to regular spiking when L-type channels are blocked (or, vice versa, the switch from spiking to complex spiking and bursting when *g*_*CaL*_ is increased) is more easily explained. The reduction in depolarizing Ca^2+^ current prevents prolonged epochs of elevated membrane potential and thus promotes regular spiking in which these are absent (Fig 5B; Fig 6B bottom two traces).

### A reduced model captures CWC dynamics

The CWC model that we have discussed so far was constructed to include the key currents characterized in past experiments. Although it produces a rich repertoire of biologically relevant dynamics, its high dimensionality complicates analysis of the mechanisms involved. Hence, we next sought to produce a reduced model that maintained much of the dynamic richness of the full one, including the experimentally observed transitions with nifedipine or iberiotoxin administration.

In the following, the term “full model” refers to the eleven-dimensional ODE model presented in system (1) and analyzed in the previous sections. The term “reduced model” indicates the reduction of the full model presented in system (23), obtained through the process presented in section Model reduction. In brief, the reduced model was obtained from the full model by removing those currents (*I*_*NaF*_, *I*_*CaT*_, *I*_*HCN*_) that we found to be relatively small throughout the dynamic regimes of the full model. Moreover, since the KCa current was found to operate near saturation, we removed the Ca^2+^ dependency from *I*_*KCa*_. Finally, all passive currents were combined into a single leak current. These changes were found to have a relatively weak effect on the model’s spiking and complex spiking behavior. The reduced model can produce the three types of activity seen in the full model – bursting, spiking, and complex spiking (Fig 7B) – although with a shift in the *I*_*App*_ values for which these occur and a slight narrowing of the range of *I*_*App*_ for which the model exhibits bursting, compared to the full model.

**Fig 7.**
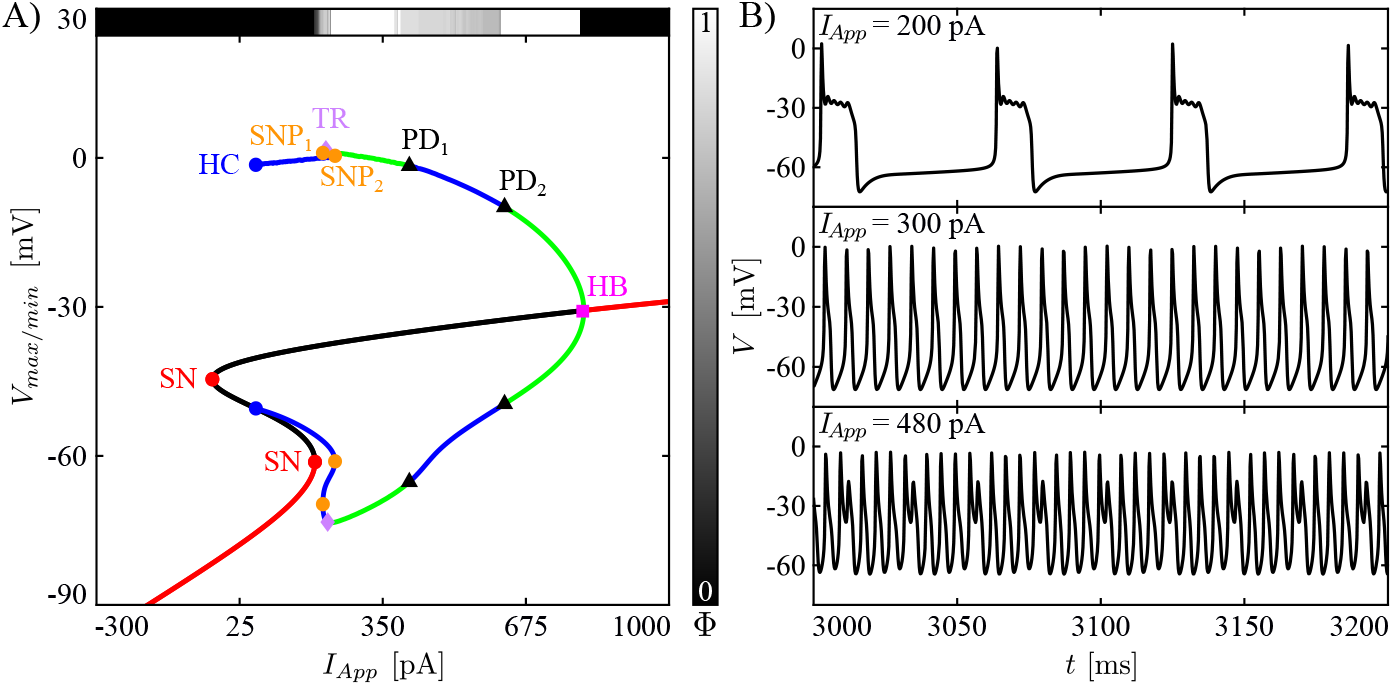
Bifurcation diagram and example voltage time series for the reduced model. (A) Bifurcation diagram of the reduced model (23) with respect to bifurcation parameter *I*_*App*_. The red/black continuous lines are stable/unstable fixed points, whereas the green/blue lines are the maximum and minimum voltages of stable/unstable periodic orbits. Other symbols are as in Fig 2. The horizontal bar on top of the BD shows how Φ changes as the *I*_*App*_ varies, whereas the vertical bar illustrates the colormap associated with the Φ values. (B) Time series of the reduced model when simulated with *I*_*App*_ equal to 200, 300 and 480 pA.

In qualitative terms, the *I*_*App*_-BD for the reduced model (Fig 7A) is mostly quite similar to that of the full model in the complex spiker regime. One exception is that the branch of stable equilibria (red part of the S-shaped curve) located at hyperpolarized voltages for *I*_*App*_ values below 196 pA no longer loses stability through a HB and rather maintains stability all the way up to a fold point (SN at approximately *V* = −60 mV). The reduced model exhibits bursting (Fig 7B, top) for *I*_*App*_ over a relatively narrow range from that at the fold point to that of a torus bifurcation (TR) where a stable family of periodic orbits, corresponding to regular spiking (Fig 7B, middle), is born. With additional increases in *I*_*App*_, regular spiking terminates in a PD bifurcation (PD_1_). The reduced model bifurcation diagram for *I*_*App*_ values above this bifurcation is simpler than that of the full model, but both produce complex spiking (Fig 7B, bottom).

### Averaging theory explains CWC dynamics at different levels of blockade of L-type Ca^2+^ and BK channels

Both the full and reduced models that we have presented feature complicated timescale structures that contribute to the details of the activity patterns that they produce, with different variables evolving on at least three disparate timescales. We observed, however, that we did not need a full dissection of the model dynamics in terms of all three timescales to understand the mechanisms involved in the reduced model dynamics that we consider. That is, within the model’s three-timescale structure, the inactivation variables of the L-type Ca^2+^ (*h*_*CaL*_) and the BK-type channels (*h*_*BK*_) act as superslow variables, evolving significantly more slowly than the other model variables. We found that we could extract fundamental insights regarding the dynamic transitions induced by varying the L-type Ca^2+^ and BK channel conductances by applying averaging theory to derive averaged dynamics for these superslow quantities, without needing to consider whether the other model variables were explicitly fast or slow.

Specifically, averaging theory is used to understand how the superslow variables change when the slow and fast components of a three-timescale system evolve along a family of attracting periodic orbits of the slow-fast subsystem. For fixed values of *I*_*App*_ and other model parameters, the regions in the (*h*_*CaL*_, *h*_*BK*_) parameter plane where the theory can be applied are therefore restricted to the sets of (*h*_*CaL*_, *h*_*BK*_) where system (24) exhibits stable periodic orbits. Accordingly, we first evaluated the bifurcation structure, the attractors, and the repellers of the slow-fast subsystem with the superslow variables *h*_*CaL*_ and *h*_*BK*_ treated as parameters. These analyses required the computation of 1P- and 2P-BDs. As described in section Averaging Theory, averaging theory is applied where stable periodic orbits arise by finding a numerical approximation of the *h*_*CaL*_ and *h*_*BK*_ superslow nullclines 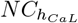 and 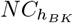 with an approach based on a periodic boundary value problem (pBVP).

Fig 8A shows a 1P-BD of the slow-fast subsystem computed using *h*_*CaL*_ as the bifurcation parameter, while fixing *h*_*BK*_ and *I*_*App*_ at 0.5 and 300 pA, respectively. In this diagram, starting from low values of *h*_*CaL*_, the BD features a branch of stable periodic orbits (green) and a branch of unstable equilibria (black), hence the slow-fast subsystem exhibits regular spiking (as shown in the simulation for *h*_*CaL*_ = 0.40 presented in the inset of the plot). As *h*_*CaL*_ increases, the branch of stable limit cycles destabilizes through a PD (PD_1_), and the latter subsequently gains stability through a subcritical HB (HB_1_). For *h*_*CaL*_ in between these two bifurcations, which occur at 0.810 and 0.902 (blue shaded region), the bifurcation diagram does not show any attractor. Simulation of the slow-fast subsystem showed that in this parameter interval, the slow-fast subsystem produces bursting (as shown in the simulation for *h*_*CaL*_ = 0.85 presented in the inset) with a number of SAOs that increases as *h*_*CaL*_ moves closer to HB_1_. A similar type of bifurcation structure has been observed in our recent work on layer V cortical neurons [27]. In that case, the mechanism for the generation of SAOs in the slow-fast subsystem was related to the existence of a folded node, but we leave the analysis of the mechanism underlying the SAOs in the CWC models for future investigations.

**Fig 8.**
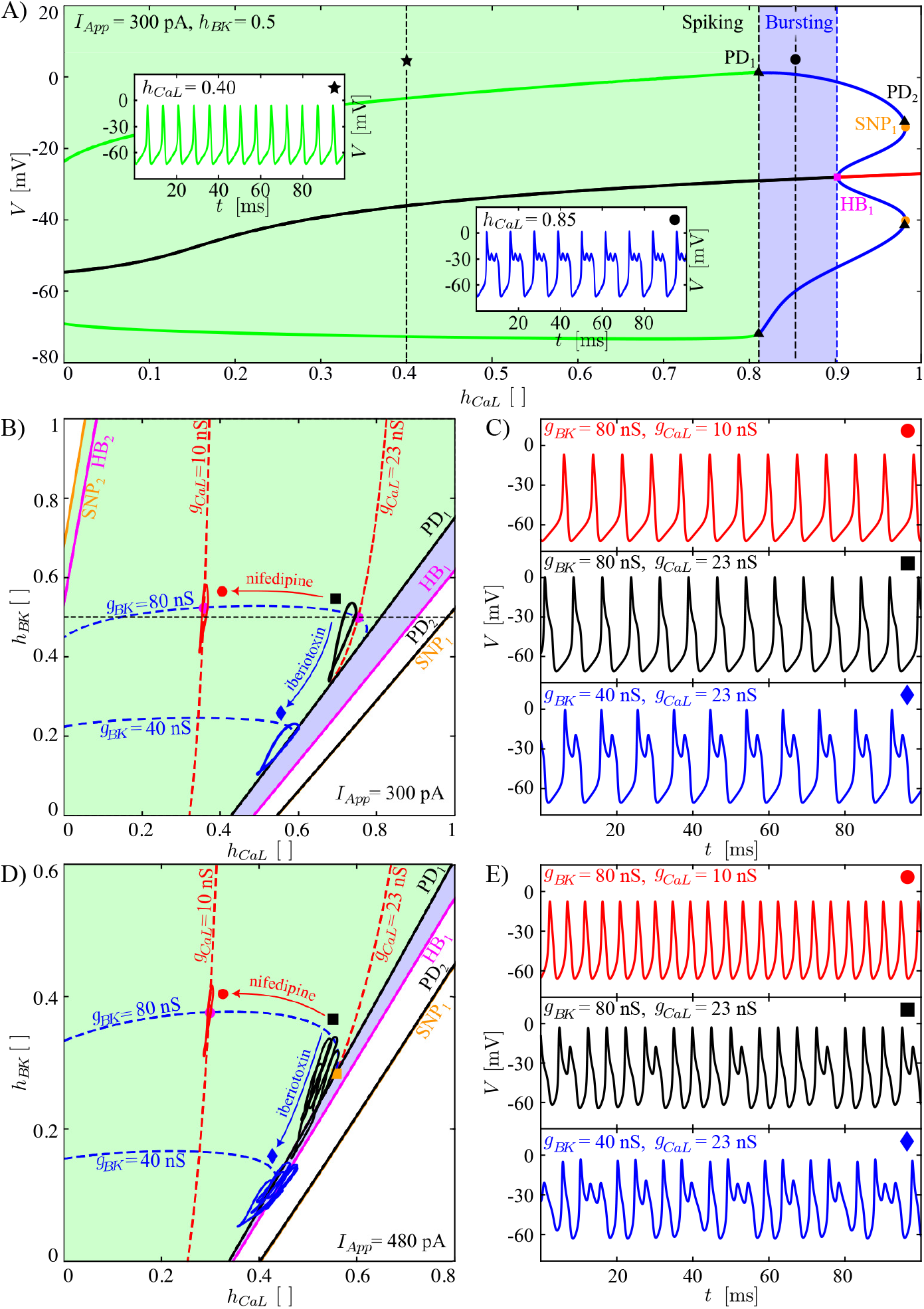
Bifurcation diagrams and time series associated with averaging theory. (A) 1P-BD of the slow-fast system 24 computed using *h*_*CaL*_ as a bifurcation parameter while fixing *h*_*BK*_ = 0.5 and *I*_*App*_ = 300 pA, respectively. The green and blue shaded regions correspond to intervals of *h*_*CaL*_ where the slow-fast subsystem exhibits regular spiking and bursting, respectively. In this panel, the inset shows a spiking (bursting) time series generated by numerically solving the slow-fast subsystem 24 for a value of *h*_*CaL*_ equal to 0.40 (0.85). (B) 2P-BD of the slow-fast system 24 computed using *h*_*CaL*_ and *h*_*BK*_ as bifurcation parameters for *I*_*App*_ = 300 pA. Bifurcation curves are labeled as in (A), but with an additional HB_2_ and SNP_2_ curves in the upper left that were not present in (A). The red and blue dashed curves are the *h*_*CaL*_ and *h*_*BK*_ averaged nullclines, corresponding to values (*h*_*CaL*_, *h*_*BK*_) where the drift of *h*_*CaL*_ and *h*_*BK*_, respectively, averaged around one cycle of the stable slow-fast periodic orbit is zero. The arrows indicate how these shift with reductions in corresponding conductances. The black square, the red circle, and the blue diamond are discussed in the text. (C) Reduced model time series for *I*_*App*_ = 300 pA, for each marker from (B). (D-E) Analogous to (B-C) but for *I*_*App*_ = 480 pA.

By following each of the bifurcation points identified in the 1P-BD based on *h*_*CaL*_ under variations of *h*_*BK*_, we next produced a 2P-BD. In the 2P-BD, the region of slow-fast periodic dynamics (Figs 8B and 8D, green shaded regions) is bounded in the direction of increasing *h*_*CaL*_ by a curve of period doubling bifurcations (Figs 8B and 8D, PD_1_ curves). We found that for *h*_*BK*_ sufficiently positive, this region is bounded below by an HB curve (Fig 8B, HB_2_), although this curve was not evident in the 1P-BD because we restricted to the physiological regime *h*_*CaL*_ ≥ 0 for *h*_*BK*_ = 0.5. Within the region of slow-fast periodic dynamics, we numerically constructed the superslow averaged nullclines (dashed blue curve for the *h*_*BK*_ nullcline and dashed red curve for the *h*_*CaL*_ nullcline). Each was computed both for a baseline condition and for a pharmacological manipulation corresponding to a reduction in the conductance of the relevant current. Inhibition of the L-type Ca^2+^ channels led to a leftward shift of the *h*_*CaL*_ superslow averaged nullcline, while inhibition of the BK channels led to a downward shift of the *h*_*BK*_ superslow averaged nullcline.

Any intersection point of the superslow averaged nullclines represents an equilibrium point of the superslow dynamics, corresponding to net zero drift of the superslow variables over one full cycle of the underlying slow-fast periodic oscillation. For baseline conductances (black square), the superslow equilibrium point is stable (Fig 8B, magenta dot). Thus, we predict that the overall reduced model system will produce regular spiking, which is exactly what we observe (Fig 8C, black square). When a moderate dose of iberiotoxin is administered (hence *g*_*BK*_ is changed from 80 to 40 nS), 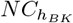 shifts downward, while 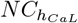 remains unchanged. In this case, as 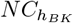 moves, the superslow stable equilibrium point slides along 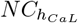, loses its stability when it crosses the fold in 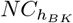, and disappears through the line PD_1_. The loss of stability of the superslow equilibrium point provides the dynamic mechanism underlying the transition from spiking to bursting in the reduced model (Figs 8B and 8C, blue diamond), with a corresponding quantative estimate for the *g*_*BK*_ value at which this event will occur. Alternatively, partial inhibition of L-type Ca^2+^ channels shifts 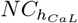 leftward, leaving 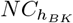 unaltered. Therefore, in this case, the super-slow equilibrium point moves leftward along the averaged *h*_*BK*_ nullcline 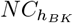 and maintains its stability, hence regular spiking persists (Figs 8B and 8C, red circle).

We repeated the same analysis for a case where the baseline dynamics is complex spiking rather than regular spiking by taking *I*_*App*_ = 480 pA (Figs 8D and 8E, black square). Here, the superslow averaged equilibrium point in the control condition exists but is unstable (orange square) and close to both the fold of 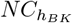 and to the line PD_1_. The instability of the equilibrium point predicts the existence of a complex form of dynamics in the reduced model. As in the previous case, reduction of the maximal BK channel conductance *g*_*BK*_ eliminates the equilibrium point (Figs 8D and 8E, blue diamond case). A new effect here is that reduction of *g*_*CaL*_ shifts the equilibrium point along 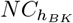 and causes it to gain stability, switching the model from complex to regular spiking (Figs 8D and 8E, red circle).

## Discussion

This work presents a biophysical model of cartwheel interneurons (CWCs), integrating information from and explaining a variety of experimental studies [11, 12, 17, 28–30]. In this paper, we focus on: (1) providing a computational demonstration of the difference between two classes of cartwheel interneurons, the spiker and the complex spiker, which confirms the experimentally motivated distinction of CWC types suggested in [12] and suggests how the two classes’ dynamics are related, and (2) elucidating the behavior of the CWCs in response to administration of iberiotoxin and nifedipine, which are pharmacological BK and L-type Ca^2+^ channel blockers [12], based on full or partial reduction of the corresponding conductances.

The model was investigated through numerical simulation and production of heatmaps (Figs 3 – 6) illustrating how the firing number [24], which reflects some of the morphological properties associated with a time series, changes under variation of model parameters *g*_*CaL*_, *g*_*BK*_, and *I*_*App*_. Unlike biological experiments, our numerical approach easily allows us to consider gradual changes in parameter values over regions of parameter space. The heatmaps highlight the relative fragility versus robustness of model dynamics as these parameters were varied, depending on the baseline regime of model activity.

Specifically, our model predicts that complex spikers can exhibit regular spiking but that this disappears when *g*_*BK*_ is slightly decreased (in response to low iberiotoxin concentrations). The exact degree of sensitivity to iberiotoxin may, of course, vary across biological interneurons; our model predicts that regular spiking will be more robust in those with higher levels of baseline BK channel conductances. Regular spikers are also predicted to exhibit complex spiking under iberiotoxin administration, but are predicted to require a significantly larger reduction in BK conductance before this change in activity occurs. Importantly, our modeling suggests that the larger baseline *g*_*K*_ and *g*_*CaL*_ and smaller baseline *g*_*NaP*_ and *g*_*K*(*Ca*)_ in the complex spiker case corresponds to a qualitative difference in bifurcation structure as *I*_*App*_ is varied, relative to the regular spiker regime. We predict that this difference will manifest in which form of dynamics will be first observed in the transition from quiescence to ongoing spiking as *I*_*App*_ is increased. In our simulations, both model types show a rapid loss of complex phenomena (i.e., complex spiking and bursting) with simulated nifedipine application. This result agrees with the observations in [12], where the administration of nifedipine significantly diminished the likelihood of generation of complex spikes.

To explain the effects of simulated pharmacological manipulations, we simplified our full model to a six-dimensional reduced model that retained the responses to current variations observed in complex spiker CWCs and preserved the dynamic transitions observed with iberotoxin and nifedipine treatment. Specifically, the reduced model exhibits bursting, spiking, and complex spiking as observed in a complex spiker CWC in response to increasing current stimulation. The reduced model is a multiple-timescale dynamical system featuring an intricate timescale structure. In this paper, we show that if we simply exploit the observation that *h*_*BK*_ and *h*_*CaL*_ evolve on the slowest (superslow) timescale, then we can capture the dynamic mechanisms underlying the transitions observed in the heatmaps between spiking and complex phenomena (i.e., bursting and complex spiking) when *g*_*BK*_ and *g*_*CaL*_ are varied. Specifically, we used averaging theory to numerically estimate the superslow nullclines in the region of (*h*_*BK*_, *h*_*CaL*_) where the slow-fast subsystem of the model, comprising its other variables, exhibits periodic oscillations. We then studied how the existence and stability of the superslow equilibrium point changes under variation of *g*_*BK*_ and *g*_*CaL*_ for two different values of *I*_*App*_ corresponding to two different baseline configurations of relevant dynamical structures. The results showed that at baseline, the superslow equilibrium point lies at a position in the (*h*_*CaL*_,*h*_*BK*_) phase space close both to a PD bifurcation curve of the reduced model slow-fast subsystem and to the fold of the *h*_*BK*_ nullcline 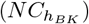, although with different stability properties in the two different baseline cases. In both, reduction of *g*_*CaL*_ shifts 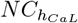 leftward, which stabilizes the equilibrium point and hence predicts that regular spiking will occur. Reduction in *g*_*BK*_ moves 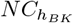 downward, promoting the destabilization of the stable equilibrium point or its disappearance through the curve of PD bifurcations. In the latter case, the theory predicts a change in the model behavior and the generation of a complex form of dynamics in the superslow variables *h*_*CaL*_ and *h*_*BK*_, corresponding to complex spiking in the overall reduced model. Consistently with the results presented in the heatmaps, averaging theory predicts which types of electrical activity will be promoted in response to an increase in *g*_*BK*_ or *g*_*CaL*_. Specifically, if *g*_*BK*_ increases, the *h*_*BK*_ nullcline will be shifted upward, promoting the stabilization of the super-slow equilibrium point and predicting the existence of a regular spiking solution. Instead, if *g*_*CaL*_ increases, the *h*_*CaL*_ nullcline will be shifted rightward, moving the super-slow equilibrium point along the 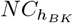, inducing a loss of stability and the emergence of complex spiking and bursting solutions.

Although our averaging analysis did exploit the multiple-timescale nature of our model, we did not pursue a more complete analysis of its timescale structure. Future studies should focus on a more detailed analysis of the various forms of complex spiking and bursting that our model produces, including the mechanisms underlying the SAOs and the transitions between bursting, spiking, and complex spiking in response to variation of *I*_*App*_, *g*_*CaL*_, and *g*_*BK*_.

To conclude, in this paper, we presented a novel biophysical ODE-based model of CWCs able to generate bursting, spiking, and complex spiking phenomena. Our analysis highlights a possible distinction between CWCs labeled as complex spikers versus spikers in previous experiments and predicts how effects of variations in an applied stimulation current and changes of BK and L-type Ca^2+^ channel conductances will modulate the electrical activity of these neurons. This work represents a first analytical step towards elucidating the mechanisms mediating the generation of complex phenomena in CWCs and how plasticity affecting L-type Ca^2+^ and BK channel conductances might influence their electrical activity and hence auditory function and dysfunction.

## Materials and Methods

### Model

We constructed a biophysical, conductance-based, whole-cell model for individual cartwheel interneurons (CWCs) using information derived from a variety of experiments.

The model equations take the form

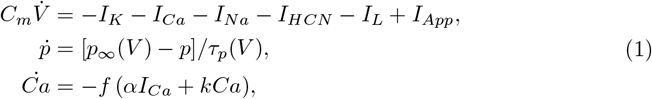

where *C*_*m*_ is the membrane capacitance, *V* is the membrane potential, *Ca* represents the calcium concentration in the intracellular compartment and, finally, *p* denotes a generic gating variable of an ion channel. Specifically, *p* ∈ {*m*_*KV*_, *m*_*KDR*_, *m*_*BK*_, *h*_*BK*_, *h*_*CaL*_, *m*_*CaT*_, *h*_*CaT*_, *h*_*NaF*_, *m*_*HCN*_ }, where the notation *m*_*X*_ indicates activation variables, while *h*_*X*_ are inactivation variables. The right-hand side of the *p* equation includes the functions *p*_*∞*_(*V*) and *τ*_*p*_(*V*), which are the steady-state and the time-scale functions, respectively. Calcium dynamics is modeled using a single compartment capturing the intracellular calcium concentration with a linear elimination rate *k*, where *f* is the ratio of free-to-total cytosolic calcium and *α* is a coefficient converting from current to molar flux. The terms *I*_*K*_, *I*_*Ca*_, and *I*_*Na*_ are the overall K^+^-, Ca^2+^- and Na^+^ currents, defined as

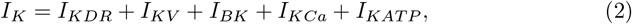

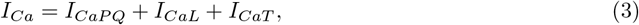

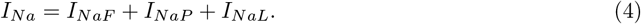

Their extended definitions are presented below and justified in the following subsections.

First, the total K^+^ current *I*_*K*_ is composed of delayed-rectifier (*I*_*KDR*_), voltage-gated (*I*_*KV*_), big-conductance (*I*_*BK*_), Ca^2+^-gated (*I*_*KCa*_) and ATP-dependent K^+^ (*I*_*KAT P*_) currents. These take the forms

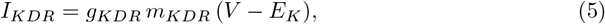

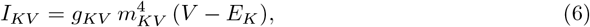

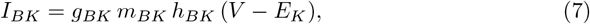

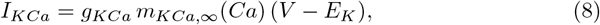

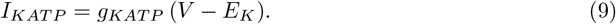

Influx of Ca^2+^ into the intracellular compartment is due to P/Q-, L-, and T-type Ca^2+^ currents (*I*_*CaP Q*_, *I*_*CaL*_, *I*_*CaT*_, respectively). The definitions of these currents are

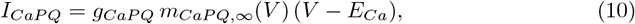

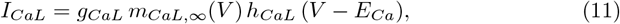

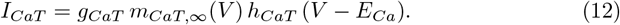

The Na^+^ currents in the model are a fast/transient type (*I*_*NaF*_), a persistent type (*I*_*NaP*_), and a Na^+^ leak current (*I*_*NaL*_), which are described by

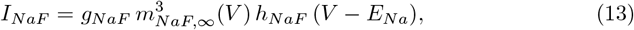

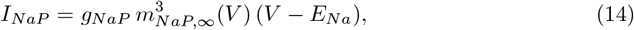

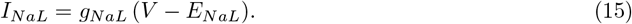

Finally, we considered three additional currents: the hyperpolarization-activated cyclic nucleotide-gated current (*I*_*HCN*_) and a general leakage current (*I*_*L*_), given by

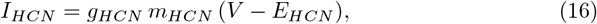

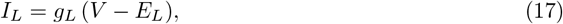

and an externally applied current denoted by the constant *I*_*App*_.

All the steady-state activation and inactivation functions implemented in the model, unless otherwise stated, take the form

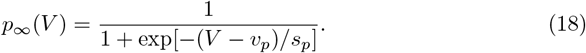

However, the steady-state activation of the Ca^2+^-gated K^+^ channel is modeled as

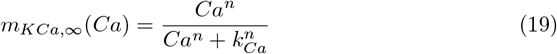

and the inactivation variable of the BK channel is modeled using a simplified formulation of the model presented in [31], where the inactivation level is controlled by the calcium concentration in the nanodomain below the mouth of P/Q Ca^2+^-channels. Hence,

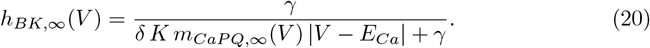

The general expression for the timescale function in (1) is

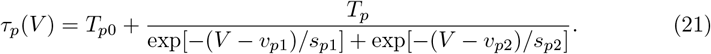

Note that the timescale function can be chosen to be modelled as voltage-independent by setting *T*_*p*_ = 0. Parameters related to the dynamics of the various gating variables are listed in Table 1. Table 2 presents the different Nernst potentials and the parameters associated with calcium dynamics, and Table 3 summarizes the values of the conductances used for a spiker and for a complex spiker CWC.

**Table 1.** Parameters used to define the kinetic properties of the ion channels involved in the model. The *STANDARD* section presents values of the parameters of the channels obeying equations (18) and (21). Those activation and inactivation variables with a voltage-independent timescale have *T*_*p*_ = 0 and *T*_*p*0_ different from zero. Those activation and inactivation variables that are assumed to be in steady-state have a ‘-’ in the columns from *T*_*p*0_ to *s*_*p*2_. The *NON STANDARD* section provides values of the remaining parameters characterizing activation of the K(Ca) channels through equation (19) and inactivation of the BK channels via equation (20).

| STANDARD |  |  |  |  |  |  |  |  |  |  |
| --- | --- | --- | --- | --- | --- | --- | --- | --- | --- | --- |
| CH. | $m/h$ | $v_p$<br>[mV] | $s_p$<br>[mV] | $T_{p0}$<br>[ms] | $T_p$<br>[ms] | $v_{p1}$<br>[mV] | $v_{p2}$<br>[mV] | $s_{p1}$<br>[mV] | $s_{p2}$<br>[mV] | REF. |
| KDR | $m$ | -11 | 8 | 0 | 13 | -100 | 50 | -39 | 65 | [29] |
| KV | $m$ | -55 | 15 | 2 | 0 | - | - | - | - | [18] |
| BK | $m$ | -22 | 8 | 2 | 0 | - | - | - | - | [17] |
| | $h$ | - | - | 5 | 0 | - | - | - | - | |
| CaPQ | $m$ | -22 | 8 | - | - | - | - | - | - | [17] |
| CaL | $m$ | -38 | 6 | - | - | - | - | - | - | [36] |
| | $h$ | -42 | -4 | 20 | 0 | - | - | - | - | |
| CaT | $m$ | -40 | 9 | 0.4 | 7 | -120 | -40 | 240 | -10 | [37] |
| | $h$ | -93 | -10 | 8 | 500 | -93 | -93 | 10 | -10 | |
| NaF | $m$ | -35 | 7 | - | - | - | - | - | - | [34] |
| | $h$ | -77 | -9 | 0 | 13 | -70 | -54 | -15 | 44 | |
| NaP | $m$ | -50 | 15 | - | - | - | - | - | - | [34] |
| HCN | $m$ | -90 | -15 | 30 | 0 | - | - | - | - | [11] |

**Table 1.** Parameters used to define the kinetic properties of the ion channels involved in the model.
| NON STANDARD |  |  |  |  |  |  |  |  |  |  |
| --- | --- | --- | --- | --- | --- | --- | --- | --- | --- | --- |
| CH. | $m/h$ | $n$<br>[ ] | $k_{Ca}$<br>[μM] | $\delta$<br>[$\frac{1}{\mu$M ms] | $K$<br>[$\frac{\mu$M}{mV}] | $\gamma$<br>[ms<sup>-1</sup>] | - | - | - | REF. |
| KCa | $m$ | 2 | 0.4 | - | - | - | - | - | - | [32, 33] |
| BK | $h$ | - | - | 0.0025 | 0.67 | 0.002 | - | - | - | [31] |

**Table 2.** This table shows the parameters of the model associated with the Nernst potentials (*E*_*Ca*_, *E*_*K*_, *E*_*L*_, *E*_*Na*_, *E*_*HCN*_ and *E*_*NaL*_), the membrane capacitance (*C*_*m*_), and the dynamics of the intracellular calcium compartment (*f, k* and *α*).

| GENERAL |  |  |  |  |  |  |  |  |
| --- | --- | --- | --- | --- | --- | --- | --- | --- |
| $E_{Ca}$ | 60 | [mV] | $E_{Na}$ | 50 | [mV] | $f$ | 0.05 | [ ] |
| $E_K$ | -85 | [mV] | $E_{HCN}$ | -20 | [mV] | $k$ | 0.3 | [ms<sup>-1</sup>] |
| $E_L$ | -40 | [mV] | $E_{NaL}$ | 0 | [mV] | $\alpha$ | 0.003 | [μM pA<sup>-1</sup> ms<sup>-1</sup>] |
| $C_m$ | 10 | [pF] | | | | | | |

**Table 3.** This table shows the conductances of a spiker (S)/complex spiker (CS) CWC model.

| CONDUCTANCES [nS] |  |  |  |  |  |  |  |  |
| --- | --- | --- | --- | --- | --- | --- | --- | --- |
| $g_X$ | CS | S | $g_X$ | CS | S | $g_X$ | CS | S |
| $g_{KDR}$ | 100 | 100 | $g_{CaPQ}$ | 5 | 5 | $g_{NaP}$ | 31 | 33 |
| $g_K$ | 32 | 29 | $g_{CaL}$ | 23 | 19 | $g_{NaF}$ | 100 | 100 |
| $g_{BK}$ | 80 | 80 | $g_{CaT}$ | 5 | 5 | $g_{NaL}$ | 1 | 1 |
| $g_{KATP}$ | 6 | 6 | $g_{KCa}$ | 10 | 11 | $g_{HCN}$ | 5 | 5 |
| $g_L$ | 0.1 | 0.1 | | | | | | |

### Potassium channels

Potassium ions move across the cellular membrane through different channels, e.g. Ca^2+^-gated K^+^ channels [12, 17, 28], Kv2 channels [29] and ATP-dependent K^+^ channels [11]. Experiments [12, 17, 28] have established the existence of big- and small-conductance K^+^ channels (BK, SK, respectively). In the model, we implement only two Ca^2+^-gated K^+^ channels, the BK and the KCa ones. The latter accounts for both the small- and the intermediate-conductance K^+^ channel contributions.

This model assumes that the BK current activates simultaneously with P/Q-type Ca^2+^ currents, hence they have the same steady-state activation function. The inactivation of the BK channels is modeled using a simplified formulation of the conductance-based model presented in [31]. The parameters *γ, K*, and *δ* are derived directly from [31]. We adjusted the steady-state activation functions for these channels and the BK maximal conductance to replicate the voltage-clamp experiments presented in Fig. 4 in [17]. The KCa channels are modeled through a Ca^2+^-dependent steady-state activation function, as presented in equation (19), derived from [32]. The value of *k*_*Ca*_ is increased to 0.4 *µ*M, which is the average of the values used in two previous models [32, 33]. The maximal conductance of the KCa current is constrained to match that of previously-studied apamine-sensitive currents [17, 28]. The properties of Kv2 channels (i.e., one fundamental component of the delayed-rectifier K^+^ class) and the dynamics of the KDR channel, specifically the *v*_*p*_ and *s*_*p*_ parameters of the steady-state activation functions of these channels and the voltage-dependent time constant, were fit to the data presented in [29]. The term *T*_*p*_ presented in this paper is scaled by a temperature factor, changing from 30 to 13 ms, which is specific for the Kv2 channels, to account for the temperature difference between the experiments in [29] and the simulations presented here. The ATP-dependent K^+^ current was modelled as a passive current. The maximal conductance *g*_*KAT P*_ was estimated to match the tolbutamide-sensitive currents detected in [11]. Specifically, in [11], the available measures distinguish between silent- and active-cartwheel when no current is applied. We aim to build a model of a silent interneuron; hence, we replicated the results associated with a silent cell. Finally, we assume the existence of an additional source of inhibitory current regulated through a generic K^+^ voltage-gated current (KV). For this purpose, we implemented in the model a K^+^ current, as presented in [18]. The maximal conductance *g*_*KV*_ was manually tuned.

### Sodium channels

Na^+^ is one of the fundamental components contributing to the generation of the electrical activity of the CWCs [12, 30]. The total Na^+^ current in these neurons is composed of TTX-sensitive and TTX-insensitive components. The former is due to persistent- and transient-type Na^+^ channels, whereas the latter reflects Na^+^ leak channels (NaLCN). The dynamics of the persistent and transient-type Na^+^ channels are taken from [34]; the maximal conductance of the persistent-Na^+^ channel *g*_*NaP*_ is constrained by the experimental data presented in [11], while that of the transient-type Na^+^ channels is manually chosen to be 3 times bigger than *g*_*NaP*_. The TTX-insensitive current, *I*_*NaL*_, is modelled as a leakage current, with a maximal conductance chosen to be of the same order of magnitude as *g*_*KAT P*_.

### Calcium channels

Kim et al. [12] found L-, T- and P/Q-types Ca^2+^ currents in CWCs. Another study made on a non-specific population of neurons of the DCN [35] found the same results. The dynamics of the P/Q-type Ca^2+^ currents in our model are adapted to meet the experiments in [17]. The dynamics of L-type Ca^2+^ currents, on the other hand, are taken from [36] and adjusted. Specifically, we changed the half-width activation of the steady-state function (*v*_*p*_) to −38 mV in our model, and we considered the inactivation timescale of the L-type calcium current to be constant and set at 20 ms. The maximal conductances of the L-, T-, and P/Q-type channels were chosen to replicate the experimental findings in [12]. Finally, the T-type Ca^2+^ currents were taken from [34, 37], which presented a model of cerebellar Purkinje interneurons.

### Hyperpolarization-activated cyclic nucleotide-gated channel

Another type of channel identified in CWCs in [11, 12] is the HCN channels. All the parameters associated with the model of this channel class were adapted to match the experiments in [11]. The dynamics of these channels are similar to those investigated in our previous work on cortical layer V neurons [27].

### Numerical methods

XPPAUT [38] was the primary tool used to solve the model system of ODEs, using the qualitative Runge-Kutta method, which is designed to preserve solution properties (e.g., positivity of gating variables), with a time step of 0.01 ms. The absolute and relative tolerances used to solve the ODE system were set to 1e-6. For the heatmap creation, we identified spikes based on crossings of a Poincaré section defined with respect to the variable *V* over a 3-second, uninterrupted simulation. MATLAB [39] was used to control the routines for the creation of heatmaps running on a computer cluster, for offline data analysis, and for figure creation. Finally, Inkscape [40] was used for image editing.

### Firing number

We computed a morphological index, the firing number [24], based on the properties of complex spiking and pseudo-plateau bursting dynamics, to quantify model activity. The voltage traces of these activity patterns include both full action potentials, or large amplitude oscillations (LAOs), and spikelets, or small amplitude oscillations (SAOs). Thanks to the existence of SAOs, these patterns resemble mixed-mode oscillations. For this reason, they can be summarized through their signature 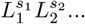 [41]. The signature of a time series can be periodic or aperiodic. In both cases, *L*_*i*_, *s*_*i*_ ∈ *ℕ* indicate the number of LAOs and SAOs, respectively, in a cycle. Therefore, for any simulated time series comprising *n* cycles, we can define 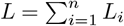 an 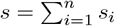, and calculate the firing number [24] as follows:

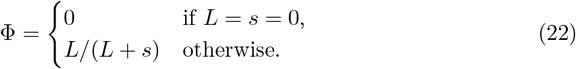

By construction, Φ ∈ [0, 1]. Specifically, Φ = 1 when the time series is comprised entirely of LAOs so that *s* = 0. Alternatively, Φ = 0 when *L* = 0, hence when the neuron exhibits SAOs only or is silent. For Φ values between 0 and 1, the time series features SAOs on top of a plateau. In this latter case, we cannot distinguish between pseudo-plateau bursting and complex spiking. However, this index can help us to discriminate between these complex patterns and regular spiking activity.

### Heatmaps

The heatmaps are constructed to go beyond the experimental results based on all-or-none channel blockage by systematically investigating how the model behaves for intermediate levels of reduction of certain transmembrane current(s) in a specific range of applied current. Partial current reductions can be simulated by changing the maximal conductance associated with a single or a set of ion channels.

We changed *I*_*App*_ and either *g*_*BK*_ or *g*_*CaL*_ over a predefined grid and, with these parameters given by each grid point, calculated the firing number Φ from the model activity. The maps were created via the following three steps:

*Step 1 - Update ICs*: For each value of the conductance (*g*_*BK*_ or *g*_*CaL*_) in the grid, we define a new initial condiction (IC) by simulating the neuron for 2 s, starting from a common, fixed baseline IC. In this pre-simulation, a constant *I*_*App*_ drives the neuron to a stable steady state, which is then used as the IC for the subsequent step. The value of the applied current taken to represent the rest condition for the model is defined to be, for the given conductance value, the largest integer multiple of −10 mV below the current 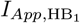 corresponding to the HB_1_ on the hyperpolarized branch of equilibria (cf. Fig 2). That is, since the location of the HB changes when *g*_*BK*_ or *g*_*CaL*_ changes, so does the value of the pre-stimulation current representing the rest condition.

*Step 2 - Model simulation*: For each value of *I*_*App*_ and *g*_*BK*_ (or *g*_*CaL*_) in the grid, we simulate the response of the neuron for 3 s, starting from the IC found as the steady-state in Step 1. We discard an initial transient of 1 s of the simulation, and save only the local maxima and minima of the remaining part of the voltage time series.

*Step 3 - Index computation*: For any combination of *I*_*App*_ and *g*_*BK*_ (or *g*_*CaL*_) in the grid, the associated series of peaks is used to calculate the firing number Φ for this grid point. More details regarding the analyses of the sequences of peaks and valleys used to calculate the index Φ are given in the Peak analysis section.

### Peak analysis

This algorithm analyzes the sequence of maxima and minima associated with a time series. It distinguishes between spiking and complex phenomena, e.g. pseudo-plateau bursting and complex spiking, and it extracts the signature of the time series to calculate the firing number Φ. The algorithm comprises the following steps:

*Step 1 - Cleaning* : The sequence of maxima and minima is cleaned from confounding phenomena, such as those peaks and valleys associated with SAOs occurring around the baseline (i.e., low-voltage, or rest state) transmembrane potential.

*Step 2 - Identification*: From the cleaned data, a series of peaks/valleys is extracted.

*Step 3 - Modality* : A test is performed to determine whether the *V*-coordinates of the valleys form a multimodal distribution. If so, the time series is associated with a complex activity pattern, and we proceed with step 4. Otherwise, the time series can be classified as regular spiking, hence Φ = 1 and the algorithm terminates.

*Step 4 - Classification*: The sequence of peaks is classified through an adaptive threshold calculated using the Otsu algorithm [42].

*Step 5 - Cropping* : The classified vector of peaks is cropped at its beginning/end so that the sequence starts/ends with a peak associated with a LAO/SAO.

*Step 6 - Calculation*: The series of maxima is converted into the sequence 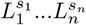. The sample is then used to compute Φ through the formulas mentioned in section Firing number.

### Model reduction

The model reduction process retains only some features of the full model. The procedure considers a model of a complex spiker and aims to produce a new model that (1) captures the transitions observed experimentally in response to administration of iberiotoxin and nifedipine and (2) maintains the transitions between bursting, spiking, and complex spiking under variation of *I*_*App*_.

The reduced model takes the form

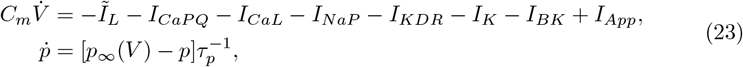

with *p* ∈ {*m*_*KDR*_, *m*_*KV*_, *m*_*BK*_, *h*_*BK*_, *h*_*CaL*_}. The values of some model parameters are those presented in Tables 1, 2 and 3, while others were modified. The form of the reduced model and the choice to alter certain parameters were based on the following reasoning:

First, we observed that the heatmaps computed by varying *I*_*App*_ and *g*_*HCN*_, *g*_*CaT*_, or *g*_*NaF*_ were not strongly sensitive to conductance variations. For this reason, despite the biological evidence for the existence of these currents in CWCs, we decided to remove these currents from the model by setting their conductances to 0 nS. This step slightly alters the *I*_*App*_ associated with the bifurcations detected in the *I*_*App*_-BD presented in Fig 2. However, with this simplification, the model still satisfies conditions (1) and (2).

In the spiking and the complex spiking conditions, the relationship between the calcium concentration and the L-type Ca^2+^ inactivation variable in the full model was linear. Moreover, for high *I*_*App*_, the KCa current was always operating at saturation, effectively acting as a leakage current. For this reason, we decided first to incorporate the KCa current into the K(ATP) current through a modification of the conductance of the latter from 6 to 17 nS. Then, we removed the calcium dynamics.

To follow the ideas presented in [43], we fixed the value of the KDR timescale at 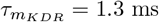, thereby obtaining a constant-*τ* model.

For the sake of compactness, we condense the leakage-type currents, *I*_*K*(*Ca*)_, *I*_*NaL*_, *I*_*K*(*AT P*)_, and *I*_*L*_, into a new leakage current (*Ĩ*_*L*_) whose conductance 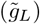 and reversal potential 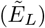 are equal to 18.1 nS and −80 mV, respectively.

These simplifications altered the model rheobase and narrowed the interval of *I*_*App*_ where the reduced model of complex spiker CWCs exhibits bursting. Moreover, the bifurcation diagram of the reduced model does not have an HB in the hyperpolarized branch of stable equilibria. This bifurcation likely disappears in a fold-Hopf bifurcation as it encounters the SN as part of the model reduction.

The reduced model obtained through these steps cannot differentiate between two different types of interneurons, spikers and complex spikers. Due to its low-dimensionality, however, it provides a useful setting in which to apply mathematical theories to explain the transitions between different activity regimes arising at different levels of inhibition of BK and L-type Ca^2+^ channels.

### Averaging Theory

Averaging theory can be used in dynamical systems with multiple timescales to understand how the slowest variables will drift along an attracting family of periodic orbits of the subsystem composed of the faster variables [25, 26, 44]. In this work, it is employed to discuss the fragility and robustness of the regular spiking presented in Figs 3 and 5.

To apply this theory, we classified the timescales of the reduced model dynamical variables. The transmembrane potential *V*, evolves with a rate that is *O*(0.1); the gating variables *m*_*KDR*_, *m*_*K*_ and *m*_*BK*_ vary with a rate that is *O*(1); the rate of *h*_*BK*_ is in between *O*(1) and *O*(10); and finally, the rate of *h*_*CaL*_ is *O*(10), with all values on a ms scale. In this scenario, we classify the variable *V* as fast, *m*_*KDR*_, *m*_*K*_ and *m*_*BK*_ as slow and finally *h*_*BK*_ and *h*_*CaL*_ as superslow. Thus, we can define the slow-fast subsystem of equation (23) as follows:

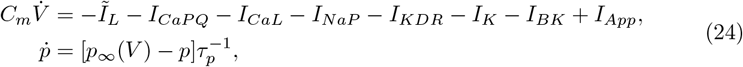

with *p* ∈ {*m*_*KDR*_, *m*_*K*_, *m*_*BK*_}. In the system presented in equation (24), the slowest variables of the reduced model, *h*_*BK*_ and *h*_*CaL*_, are considered as parameters.

The averaging theory is applied by numerically reconstructing the superslow averaged nullclines and studying the stability of any superslow equilibrium points where these nullclines intersect. In the following, the procedure to compute the *h*_*BK*_ superslow nullcline 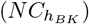 is presented. A similar approach has been adopted for the calculation of the *h*_*CaL*_ superslow average nullcline 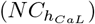. We indicate with *γ* a stable limit cycle existing for a given combination of the parameters *h*_*CaL*_ and *h*_*BK*_ in the slow-fast subsystem presented in equation (24). We define the superslow average variation along *γ* as

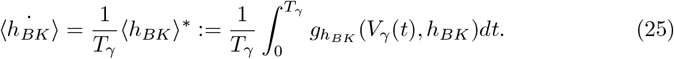

The function 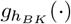 corresponds to the right-hand side of the *h*_*BK*_ differential equation in the reduced model. *V*_*γ*_(*t*) indicates the temporal evolution of the *V*-coordinate of *γ*, while *T*_*γ*_ is the period of *γ*. On the right hand side of equation (25), *h*_*BK*_ is required to take values where a limit cycle *γ* exists. The quantity 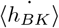 represents the average variation along *γ* of the superslow variable *h*_*BK*_. The *h*_*BK*_ superslow average nullcline is

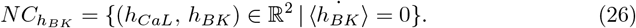

To compute this set of points, we adapted an approach based on continuation of solutions to a periodic boundary value problem (pBVP) presented in [44]. We rescale *t* in equation (24) by *T*_*γ*_, hence 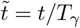, and we extend the system with three additional dynamical variables, namely ⟨*h*_*BK*_⟩, *h*_*BK*_, and *T*_*γ*_, as follows:

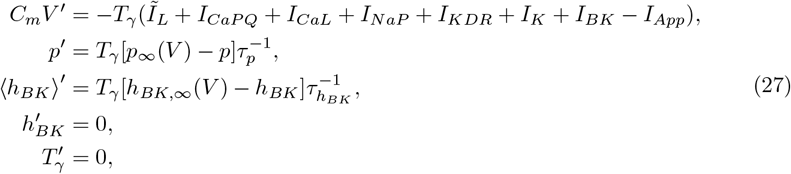

where the prime symbol denotes differentiation with respect to the new time variable 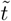. The last two equations in (27) act as free parameters in the numerical continuation.

The limit cycle *γ* is still a solution of the extended system in equation (27) but now, with respect to the new temporal variable 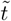, the period is unitary. The system in equation (27) is solved using the periodic, boundary, and initial conditions

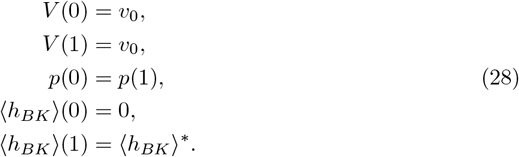

The pBVP (27), (28) can be solved only if the dynamical variables and the parameter *v*_0_ are properly initialized. ⟨*h*_*BK*_⟩^*∗*^ is the value computed through the integral in equation (25) along *γ*. The value *v*_0_ must be set to the value of *V* assumed along *γ* at the same instant of time chosen to initialize the periodic conditions set for the dynamical variables *p*.

The construction of 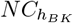 is completed through the following four steps:

*Step 1 - 1P-BD:* The one-parameter bifurcation diagram (1P-BD) of the slow-fast subsystem in (24) is computed using *h*_*BK*_ as the bifurcation parameter by fixing *h*_*CaL*_ to 0. We performed this step using XPPAUT [38], which we also used to calculate an approximation of all the stable limit cycles in the branch of detected stable periodics (NPR is set to 1).

*Step 2 - Initialization:* The resulting BD is loaded in MATLAB with XPPLORE [45] to compute ⟨*h*_*BK*_⟩^*∗*^ for every stable limit cycle saved from the generation of the 1P-BD. After this step, the limit cycle with the lowest |⟨*h*_*BK*_⟩|^*∗*^ is selected and the values of *h*_*CaL*_, *h*_*BK*_, *T*_*γ*_ and ⟨*h*_*BK*_⟩^*∗*^ are extracted to initialize the *ode* file containing an implementation of system (27) together with the conditions in (28).

*Step 3 - Continuation:* First, AUTO in the XPPAUT package is used to find a *γ* value where ⟨*h*_*BK*_⟩(1) is zero through a numerical continuation along the ⟨*h*_*BK*_⟩^*∗*^ parameter. Second, the solution to the pBVP is continued along *h*_*CaL*_. This last step calculates 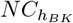.

*Step 4 - Cropping:* In the last step, the outcome of the previous numerical continuation is loaded in MATLAB through XPPLORE and the curve is cropped in the region of the parameter space (*h*_*CaL*_, *h*_*BK*_) where the slow-fast subsystem in (24) admits stable limit cycles.

The same method is iterated to calculate the super-slow nullcline 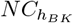 and 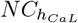 for different values of *I*_*App*_. Finally, the super-slow equilibrium point is obtained as the intersection between 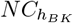 and 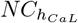, estimated numerically. Its stability is investigated through visual inspection of the flow around the point based on numerical simulations.

